# POIROT: A powerful test for parent-of-origin effects in unrelated samples leveraging multiple phenotypes

**DOI:** 10.1101/2022.11.28.517712

**Authors:** S. Taylor Head, Elizabeth J. Leslie, David J. Cutler, Michael P. Epstein

## Abstract

**Motivation:** There is widespread interest in identifying genetic variants that exhibit parent-of-origin effects (POEs) wherein the effect of an allele on phenotype expression depends on its parental origin. POEs can arise from different phenomena including genomic imprinting and have been documented for many complex traits. Traditional tests for POEs require family data to determine parental origins of transmitted alleles. As most genome-wide association studies (GWAS) instead sample unrelated individuals (where allelic parental origin is unknown), the study of POEs in such datasets requires sophisticated statistical methods that exploit genetic patterns we anticipate observing when POEs exist. We propose a method to improve discovery of POE variants in large-scale GWAS samples that leverages potential pleiotropy among multiple correlated traits often collected in such studies. Our method compares the phenotypic covariance matrix of heterozygotes to homozygotes based on a Robust Omnibus Test. We refer to our method as the Parent of Origin Inference using Robust Omnibus Test (POIROT) of multiple quantitative traits.

**Results:** Through simulation studies, we compared POIROT to a competing univariate variance-based method which considers separate analysis of each phenotype. We observed POIROT to be well-calibrated with improved power to detect POEs compared to univariate methods. POIROT is robust to non-normality of phenotypes and can easily adjust for population stratification and other confounders. Finally, we applied POIROT to a GWAS of quantitative anthropometric measures at birth. We identified two loci of suggestive significance for follow-up investigation.

## 1 INTRODUCTION

Most genome-wide association studies (GWAS) implicitly assume the magnitude and direction of effect of a genetic variant on expression of a phenotype is independent of whether the variant was maternally or paternally inherited. However, there exists a distinct class of genetic variants for which this assumption is violated. Such variants harbor a parent-of-origin effect (POE) wherein the effect of an allele on a trait depends on whether it was transmitted from the mother or the father (Lawson *et al*., 2013). A substantial proportion of the variation in complex traits is not explained by the additive effects of common single nucleotide polymorphisms (SNPs) across the genome. POEs may represent an important contribution to this missing heritability (Guilmatre and Sharp, 2012).

There are multiple cited biological mechanisms by which POEs can arise in mammals. These include maternal intrauterine environment effects and effects of the maternal mitochondrial genome. However, the most frequently considered mechanism is genomic imprinting (Rampersaud *et al*., 2008). This epigenetic phenomenon was formally discovered in the 1980s primarily through embryological experiments (Reik and Walter, 2001). In imprinting, the maternal and paternal alleles undergo differential epigenetic modifications that leads to parent-of-origin-specific transcription of the gene copies. Many imprinted genes tend to be found in clusters across the genome. Regulation of the expression of these clustered genes is under control of an imprinting control region (ICR), the mechanisms of which are complex (Barlow, 2011). These ICR are often characterized by repetitive sequences and located near imprinted genes. It is estimated that only approximately 1% of mammalian genes are subject to imprinting. However, there has been growing evidence for the existence of POE variants for a wide range of hereditary traits (Peters, 2014). Classic examples of POE-associated diseases include Prader-Willi syndrome and Angelman syndrome. These diseases result from imprinted genes at 15q11-15q13 when only maternal or paternal copies are expressed, respectively (Aypar *et al*., 2014). Considerable research has further suggested POEs originate for a wide spectrum of complex traits, including obesity-related phenotypes, type 2 diabetes, basal-cell carcinoma, attention-deficit/hyperactivity disorder, schizophrenia, and breast cancer (Rampersaud *et al*., 2008; Giannoukakis *et al*., 1993; Temple *et al*., 1995; Huxtable *et al*., 2000; Polychronakos and Kukuvitis, 2002; Hoggart *et al*., 2014; Dong *et al*., 2005; Kong *et al*., 2009; Wang *et al*., 2012; Palmer *et al*., 2006).

To detect variants demonstrating POEs, studies have historically required genotype data from related individuals to ascertain parental ancestry of the inherited alleles. In the case of available parent-offspring trio or other forms of familial genomes, there are well-established methods to detect POEs (Connolly and Heron, 2015; Weinberg *et al*., 1998; Cordell *et al*., 2004; Howey and Cordell, 2012; Ainsworth *et al*., 2011; Sinsheimer *et al*., 2003; Howey *et al*., 2015; Becker *et al*., 2006; Zhou *et al*., 2012; Weinberg, 1999). These approaches often test for a mean difference in allele effect based on grouping offspring by parent-of-origin of the allele. These mean-based tests are intuitive and optimally powered given sample size. Yet, the requirement of trio or more general family data severely limits this sample size in practice. This, in consequence, limits genome-wide discovery of the modest genetic effects that we anticipate for complex human traits.

Rather than rely on family studies of limited sample size to detect POEs, researchers have recently transitioned to detecting the phenomenon in GWAS-scale cohorts. This practice requires innovative statistical methods to deal with missing parental ancestry information. For example, Kong et al. inferred parental origin of alleles when parental genotype data are not available by first phasing Icelandic probands. For each of the proband haplotypes, they searched a genealogy database for the closest typed maternal and paternal relatives carrying that haplotype (Kong *et al*., 2009). For each haplotype, they constructed a robust score comparing the meiotic distances between the proband and these two relatives to quantify the likelihood of maternal or paternal transmission and ultimately assign parental origin. While this approach solves the issue of small sample sizes, power is still impacted by the potential inaccuracy or uncertainty in haplotypic reconstruction.

More recently, Hoggart et al. described a novel statistical method for detecting POEs for a single quantitative trait using GWAS data of unrelated individuals (Hoggart *et al*., 2014). The authors illustrated that the existence of a POE results in increased phenotypic variance among heterozygotes compared to homozygotes. They tested for this variance inflation using a robust version of the Brown-Forsythe test. The method successfully identified previously undocumented POE associations of two SNPs with body mass index (BMI). This work has enabled POE analysis in population studies of biobank scale. However, such variance-based tests are often underpowered compared to their corresponding mean-based tests described above when allelic parental origin is known (Struchalin *et al*., 2010). Furthermore, the method only tests for parent-of-origin-dependent associations between a genetic variant and a single phenotype.

A sizable proportion of genes in the GWAS catalog are pleiotropic (Chesmore *et al*., 2018). These genes affect more than one biological process, in turn associating with multiple (correlated) phenotypes (He and Zhang, 2006). Analyzing the joint effects of a gene on more than one trait can often result in a marked increase in power over univariate approaches (Kocarnik and Fullerton, 2014; Solovieff *et al*., 2013; O’Reilly *et al*., 2012). Importantly, well-established POEs in humans are usually the result of embryonic silencing of one parental allele. As this silencing generally occurs early in development, its effects are likely to present in all or nearly all tissues expressing the gene. When differential silencing of this gene affects multiple tissues, this can yield POEs for several distinct phenotypes. Joint analysis of multiple traits can leverage this potential pleiotropy to improve power over univariate variance-based POE tests while simultaneously reducing multiple testing burden of multiple phenotypes.

Here, we expand on the concept initially suggested by Hoggart et al. to develop a test for POEs in genetic studies of unrelated individuals that considers multiple quantitative phenotypes. We show that a pleiotropic POE variant will not only induce differences in the variance of POE traits between heterozygotes and homozygotes, but also in their corresponding covariances. In our method, POIROT (Parent-of-Origin Inference using Robust Omnibus Test), we test for equality of phenotypic covariances matrices between heterozygous and homozygous groups. Specifically, we use the robust omnibus (R-Omnibus) test (O’Brien, 1992) to accommodate highly skewed traits. We first provide background on the statistical construction of our test statistic using the R-Omnibus framework. Next, through simulations, we demonstrate that our proposed method properly controls type I error and achieves superior power compared to the univariate approach of Hoggart et al. We apply our method to GWAS data of fetal growth phenotypes from the Hyperglycemia and Adverse Pregnancy Outcome (HAPO) study and identify two potential POE loci. We conclude with a discussion of our findings and proposed research to extend this work.

## 2 METHODS

### 2.1 Phenotype Model

Using the notation of Hoggart et al., consider one biallelic SNP with reference allele A and alternative allele B (Hoggart *et al*., 2014). Assume we have collected *n*_*AA*_ individuals who have the homozygous AA genotype, *n*_*BB*_ individuals who have the homozygous BB genotype, and n_AB_ individuals who are heterozygous. Further assume we have collected *K* > 1 continuous phenotypes on all subjects and that we have already adjusted these phenotypes for the effects of non-genetic confounders like principal components of ancestry.

We first model phenotypes in homozygous AA subjects. Let 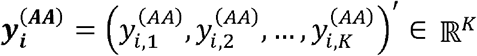 be the vector of phenotypes for the *i*^th^ AA individual. We can represent 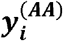 using the following framework

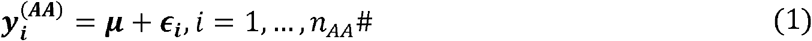

Within (1), ***µ*** = (*µ*_1_, …, *µ*_*K*_)′ is the *K* × 1 vector of phenotype means in AA subjects and ***ϵ***_*i*_ (*ϵ*_*i*1_, …, *ϵ*_*iK*_)′ is the *K* × 1 vector of error terms. We assume that E[***ϵ***_***i***_] = **0**_***K***_ and Cov[***ϵ***_***i***_] = **Σ**, where **Σ** is the *K* × *K* variance-covariance matrix of the vector of error terms.

We next model phenotypes in homozygous BB subjects. Let 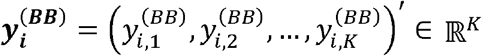 be the vector of phenotypes for the *i*^th^ BB individual. Further, let *β*_*Mk*_ and *β*_*Pk*_ represent the effect of the maternally-inherited and paternally-inherited B allele, respectively, on the *k*th phenotype. If there is no association between this SNP and the *k*th phenotype, it follows that *β*_*Mk*_ = *β*_*Pk*_ = 0. If there is a marginal association between this SNP and the *k*th phenotype, but there is no POE present, then *β*_*Mk*_ = *β*_*Pk*_ ≠ 0. With this notation defined, we can model 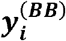 as

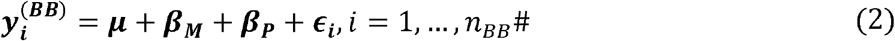

Where ***µ*** is as defined previously for (1), ***β***_***M***_ (*β*_*M*l_, …, *β*_*MK*_)′ is the *K* × 1 vector of maternal effects of the B allele on each of the *k* phenotypes, and ***β***_***P***_ (*β*_*M*l_, …, *β*_*MK*_)′ is the *K* × 1 vector of corresponding paternal effects of the B allele. Each element of ***β***_***M***_ and ***β***_***P***_ is assumed to be a fixed effect. Just as for the AA subjects in (1), we assume that E[***ϵ***_***i***_] = **0**_***K***_ and Cov[***ϵ***_***i***_] = **Σ**.

Lastly, we consider heterozygous AB individuals who carry only one copy of the alternative allele B. Let 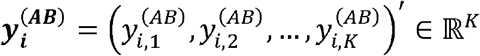 be the vector of phenotypes for the *i*th heterozygote. We can model this vector as

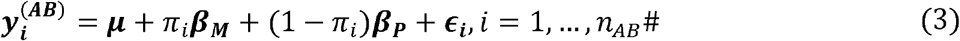

In (3), *π*_*i*_ is an indicator random variable where *π*_*i*_ = 1 if individual *i* received the B allele from the mother and *π*_*i*_ = 0 if individual *i* received the B allele from the father. We assume *π*_*i*_ ∼ Bernoulli(½), as we expect that half of heterozygotes will have maternally-derived B alleles. The maternal and paternal effect vectors are as defined as for the model of BB subjects. We also assume that E[***ϵ***_***i***_] = **0**_***K***_ and Cov[***ϵ***_***i***_] = **Σ** In other words, the covariance matrix of the error terms is the same within all three genotype groups.

Based on the derivations above, we can calculate the phenotype covariance matrix for each genotype category. Based on equations (1) and (2), it is straightforward to show that the phenotype covariance matrix of AA individuals (**Σ**) is the same as the analogous matrix for BB individuals. Therefore, we can define **Σ**_***Hom***_ = **Σ** as the phenotypic covariance matrix for all homozygous subjects. For heterozygous AB subjects modeled in equation (3), we can show that (assuming *π*_*i*_ ⊥ *ϵ*_*i*_ ∀ *i,i* ∈ (1,…, *n*_*AB*_) the phenotype covariance matrix for heterozygotes is 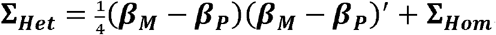. Defining *b*_*k*_ = *β*_*Mk*_ − *β*_*Pk*_ (*k* = 1, …,*K*), we can show that **Σ**_***Het***_ **= Σ**_***Hom***_ if and only if

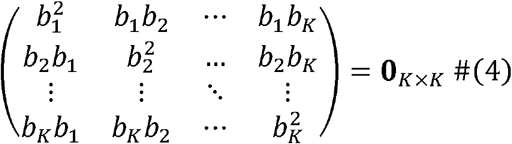

This observation motivates the use of a test of equality of two covariance matrices for detecting POEs in a population-based sample where we cannot explicitly observe *π*_*i*_. If a POE SNP exists for any phenotype *k*, then *b*_*k*_ ≠ 0 and 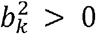. Thus, the *k*th diagonal element of **Σ**_***Het***_ will be larger than the corresponding element of **Σ**_***Hom***_. Furthermore, if the SNP has POEs on two phenotypes *k* and *k*’, then *b*_*k*_*b*_*k*′_ ≠ 0 The *kk*’ element of **Σ**_***Het***_ will also be different from the corresponding off-diagonal element of **Σ**_***Hom***_.

### 2.2 POIROT Method to Detect POE SNPs

We can test the null hypothesis that no POEs exist at a given SNP for any of the *K* phenotypes under study (H_0_: **β**_***M***_ ***=* β**_***P***_) by equivalently testing H_0_: **Σ**_***Het***_ **= Σ**_***Hom***_. In our proposed method POIROT, we test for equality of these phenotypic covariance matrices between homozygotes and heterozygotes using the robust omnibus (R-Omnibus) test of O’Brien (O’Brien, 1992). POIROT uses R-Omnibus rather than the traditional Box’s M test (Box, 1949) to test covariance differences since the latter is highly sensitive to deviations of phenotypes from multivariate normality. This can lead to a undesirable elevation in type I error rates (Tiku and Balakrishnan, 1984).

To derive the R-Omnibus test, we first center the phenotypes by the median within each genotype group (AA, AB, BB). This step is necessary if a marginal association exists between the alternative allele and a given phenotype. In that event, the variance of original phenotype values among aggregate homozygous subjects (AA, BB) would be erroneously inflated. We next group these centered phenotypes by homozygote versus heterozygote status. Let 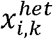 be the *k*th centered phenotype of the *i*th heterozygote (*i* = 1, …, *n*_*AB*_) and 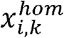 be the *k*th phenotype of the *i*th homozygous (AA and BB combined) individual (*i* = 1, …, *n*_*AA*_ + *n*_*BB*_). We then calculate the median of each phenotype *k* in heterozygotes and homozygotes separately. Let 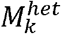 be the median of the *k*th phenotype in the *n*_*AB*_ heterozygotes. Correspondingly, let 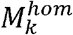 be the median of the *k*th phenotype in the *n*_*AA*_ + *n*_*BB*_ homozygotes. For heterozygotes and homozygotes separately, we then calculate for phenotypes *k* and *k’*:

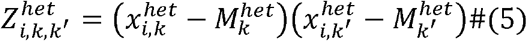

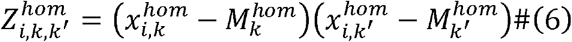

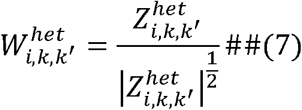

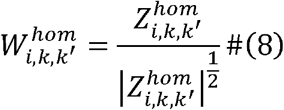

In (7) and (8), we standardize the *Z* measures by dividing by the square root of their absolute values. We consider 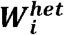 to be the vector of *W* values for the *i*th heterozygous subject, and 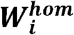 is the corresponding vector of *W* values for the *i*th homozygous subject. We then perform a two-sample Hotelling’s T^2^ test (Hotelling, 1931) comparing our two sets of *p* = (*K*^2^ + *K*)/2 sample means 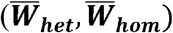. There are *p* dependent variables being compared between heterozygotes and homozygotes as this corresponds to the number of upper-triangular elements in the phenotypic covariance matrix. We calculate the test statistic 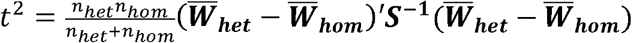, where ***S***^**−1**^ is the inverse of the pooled covariance matrix estimate. Under the null, our test statistic *t*^2^∼*T*^2^ (*p, n*_*het*_ + *n*_*hom*_ − 2) (Hotelling, 1931). The test can also be viewed as a one-way multivariate analysis of variance test (MANOVA).

### 2.3 Simulation Study

We conducted a variety of simulation studies to determine POIROT’s ability to detect POEs while maintaining proper rates of type I error. We considered *K* = 3, 6, or 10 phenotypes and *n* = 3,000, 5,000, or 10,000 unrelated individuals. To generate phenotypes for each round of simulation, we first randomly generate *K* intercepts from a standard normal distribution to form the *K* × 1 vector ***μ***. This corresponds to the mean vector of phenotypes among AA homozygotes. For simplicity, we assume the diagonal elements of the matrix **∑**, corresponding to the variances of the random error terms, are all equal to one. We assume the *K* phenotypes exhibit one of three possible levels of pairwise correlation (low, medium, or high). We assume the pairwise trait correlations are randomly drawn from a uniform distribution. To simulate phenotypes exhibiting “low” correlation, we assume this is a Uniform(0,0.3) distribution. For phenotypes of “medium” and “high” correlation, we assume a Uniform(0.3,0.5) and Uniform(0.5,0.7) distribution, respectively. These random draws are used to populate the off-diagonal elements of **∑**.

Once we have constructed **∑**, we then randomly generate *n* maternal and paternal genotypes for a given SNP by sampling twice from a Bernoulli(*p* = MAF [minor allele frequency]) for each parent. To generate offspring genotypes, we sample from a Bernoulli(*p* = 0.5) distribution to determine which maternal allele and which paternal allele is transmitted. Thus, we can now assign all *n* offspring to one of four genotype groups: (1) AB with maternal reference/paternal alternative, (2) AB with paternal reference/maternal alternative, (3) AA, and (4) BB. We then simulate the phenotypic error vector for all *n* unrelated offspring by drawing from a multivariate distribution with mean 0 and variance-covariance matrix **Σ**. The respective fixed *K* × 1 maternal and paternal effect vectors of the alternative allele (***β***_***M***_, ***β***_***P***_) are constructed depending on the specific null or alternative scenario under consideration. We then add these vectors to the random error and intercept term in concordance with the genotype group of each individual, as described in Section 2.1.

For type I error rate simulations, as described above, we assume these phenotypes have pairwise-trait correlation of levels low, medium, or high. To reflect the scenario where there exist no POEs or marginal effects of the alternative allele at the locus for any phenotype, we assume that ***β***_***M***_ = ***β***_***P***_ = **0**. We also considered a second null scenario wherein a marginal association exists for the variant that is not specific to the parent of origin, i.e., ***β***_***M***_ = ***β***_***p***_ ≠ **0**. However, we note that if the same seeds are used in simulating the data, this marginal fixed effect is effectively removed when centering phenotypes by genotype group. The resulting test statistics are equivalent to the first null scenario. We first consider the circumstance where the random error terms are drawn from a normal distribution, i.e., the error follows *MVN*_*K*_ (**0, ∑**) and assume a MAF of 0.25. For each of the 27 combinations of number of phenotypes, sample size, and pairwise-trait correlation, we conducted 50,000 null simulations. To evaluate the robustness of our method to highly skewed phenotypes, we then repeated these parameter settings with non-normal random error terms. In particular, we utilize the method of Vale and Maurelli to simulate multivariate non-normal error terms assuming a skewness of two and excess kurtosis of two for each phenotype (Vale and Maurelli, 1983). An example distribution of such a phenotype is illustrated in Supplemental Figure 1.

Next, we investigated the power of our test when POEs do in fact exist for a locus. We again considered K = 3, 6, or 10 normally distributed phenotypes. We assumed 1, 2, or 3 had parent-of-origin specific associations with the variant. When the number of affected phenotypes is greater than one, this corresponds to pleiotropy. For these scenarios, we assumed ***β***_***P***_ = **0** and *β*_*Mk*_ **=** 0.5, 0.6, or 0.75 for each phenotype *k* harboring a POE. All other elements of the maternal effect vector are 0 for the phenotypes with no POE associations. We again considered low, medium, and high pairwise-trait correlations. We assumed a MAF of 0.25 and sample sizes of 5,000, and 10,000. We applied our method to 5,000 simulated datasets for each of the 162 settings and calculated power at significance level *α* ∈ {0.005, 5 × 10^−4^}. We also compared the performance of POIROT to the corresponding univariate test of Hoggart et al. (Hoggart *et al*., 2014). For the univariate test, we first calculated power using standard Bonferroni correction. Power was calculated as the proportion of loci for which the minimum observed p-value across the *K* phenotypes tested was less than *α*/*K*. Given that these phenotypes are correlated and therefore may not reflect *K* independent tests, this approach can be overly conservative. Thus, we implemented a second more liberal approach that estimates the true number of independent tests, *K*_*eff*_, which corresponds to the minimum number of principal components (PCs) explaining 90% of the variation in our *K* phenotypes. We then calculated power of the univariate approach as the proportion of loci for which the minimum observed p-value was less than *α*/*K*_*eff*_ (Gao *et al*., 2008; Broadaway *et al*., 2016). We then repeated these parameter settings for assessing power of POIROT with non-normal phenotypes, as described for null simulations.

### 2.4 Application of POIROT to HAPO Study

Moore and Haig hypothesized that genomic imprinting is a result of the opposing interests of the maternal and paternal genomes on fetal development (Moore and Haig, 1991). In particular, the paternal genes favor greater nutrient transfer from mother to embryo to make offspring larger and thus more likely to survive. However, larger offspring represent a greater challenge to the mother in terms of the ability of the offspring to safely fit through the birth canal and a potential threat to future reproductive success. This can lead maternal genes to favor a more modest nutrient transfer to the embryo. Based on this evolutionary theory, anthropomorphic phenotypes at birth like total weight or head circumference carry high potential to be imprinted and are likely candidates for potential POEs. Therefore, to assess the utility of POIROT for detecting POEs on continuous phenotypes using published population-based GWAS data, we utilized genotype and phenotype data from the Hyperglycemia and Adverse Pregnancy Outcome Study (HAPO Study Cooperative Research Group, 2009; HAPO Study Cooperative Research Group *et al*., 2008, 2006; HAPO Study Cooperative Research Group, 2002). This study explored genetic variation associated with offspring size measures at birth, maternal glucose tolerance indicators, and the interaction of maternal/fetal genetic and environmental factors on these phenotypes using paired maternal and offspring DNA.

Through dbGaP (accession number phs000096.v4.p1), we obtained data on six quantitative phenotypes related to infant size at birth (birth weight, birth length, head circumference, flank skinfold thickness, subscapular skinfold thickness, triceps skinfold thickness). Relevant covariates included PCs, infant sex, gestational age at birth, maternal pre-pregnancy BMI, and maternal smoking status during pregnancy (none, 1-10 per day, >10 per day). While this is a multi-ethnic study, we restricted our analysis to infants of European ancestry. Subjects were genotyped using the Illumina Human610 Quad BeadChip. Prior to lift over and imputation, we excluded infants with genotype missingness greater than 10%, variants with missingness greater than 2%, variants with MAF < 0.005, and variants with Hardy-Weinberg Equilibrium *p* < 1e-8. We then lifted over genotype array data to hg38 and followed the pre-imputation quality control pipeline provided at https://www.well.ox.ac.uk/∼wrayner/tools/#Checking. We performed imputation using the TOPMed Imputation Server (reference panel TOPMed Freeze 5) (Taliun *et al*., 2021; Das *et al*., 2016; Fuchsberger *et al*., 2015). We kept only those variants with Rsq > 0.3. After quality control and imputation, 6,219,272 SNPs with MAF > 0.05 remained for analysis across 1,289 unrelated infants. All mothers indicated no illicit drug use during pregnancy. Covariate adjustment was performed by first fitting a linear model for each phenotype and extracting the residuals as the new adjusted phenotypes. We then applied POIROT to these six adjusted phenotypes to jointly test for POEs across the genome. We compared the findings of our approach to those from the method of Hoggart et al. performed on each phenotype individually.

## 3 RESULTS

### 3.1 Type I Error Rate

We summarize the type I error of null scenarios with a sample size of 5,000 individuals using Quantile-Quantile (QQ) plots in Figure 1 (normal traits) and Figure 2 (non-normal traits). Across the settings considered, our method yields the expected distribution of p-values under the null hypothesis of no POEs for any single phenotype. The distribution of the *p*-values is again as expected under the null when we have non-normality of phenotypes (Figure 2), suggesting our method remains robust. We summarize the empirical type I error rates of our proposed test and the competing univariate approach at significance level *α* ∈ {0.05, 0.005, 5 X 10^−4^,5 × 10^−5^} in Supplemental Table 1. POIROT maintained appropriate type I error across all scenarios for normally distributed traits. We observed slightly higher error when 6 or 10 highly-skewed non-normal phenotypes were tested. The univariate approach with correction for *K*_*eff*_ tests showed minor inflation with 6 or 10 highly correlated phenotypes.

**Figure 1.**
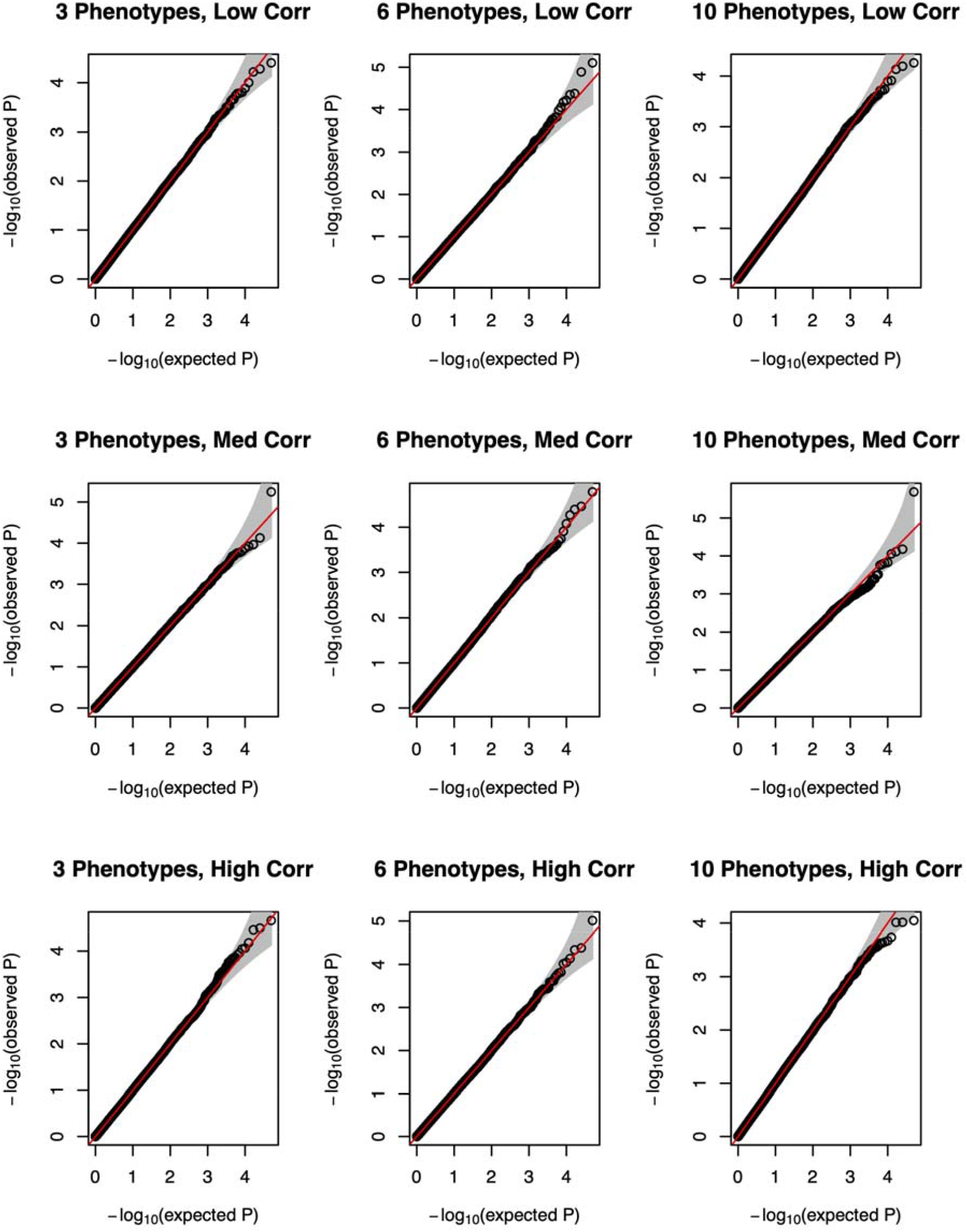
QQ plots of p-values for proposed parent-of-origin effect test under the null hypothesis ***β***_***M***_ = ***β***_***P***_ = **0** using a series of 50,000 simulations of 5,000 individuals using 3 (left column), 6 (middle column) or 10 (right column) continuous normal phenotypes. MAF is assumed to be 0.25. Horizontal panels depict level of pairwise-trait correlation (low, medium, high). Abbreviations: QQ, quantile-quantile; MAF, minor allele frequency.

**Figure 2.**
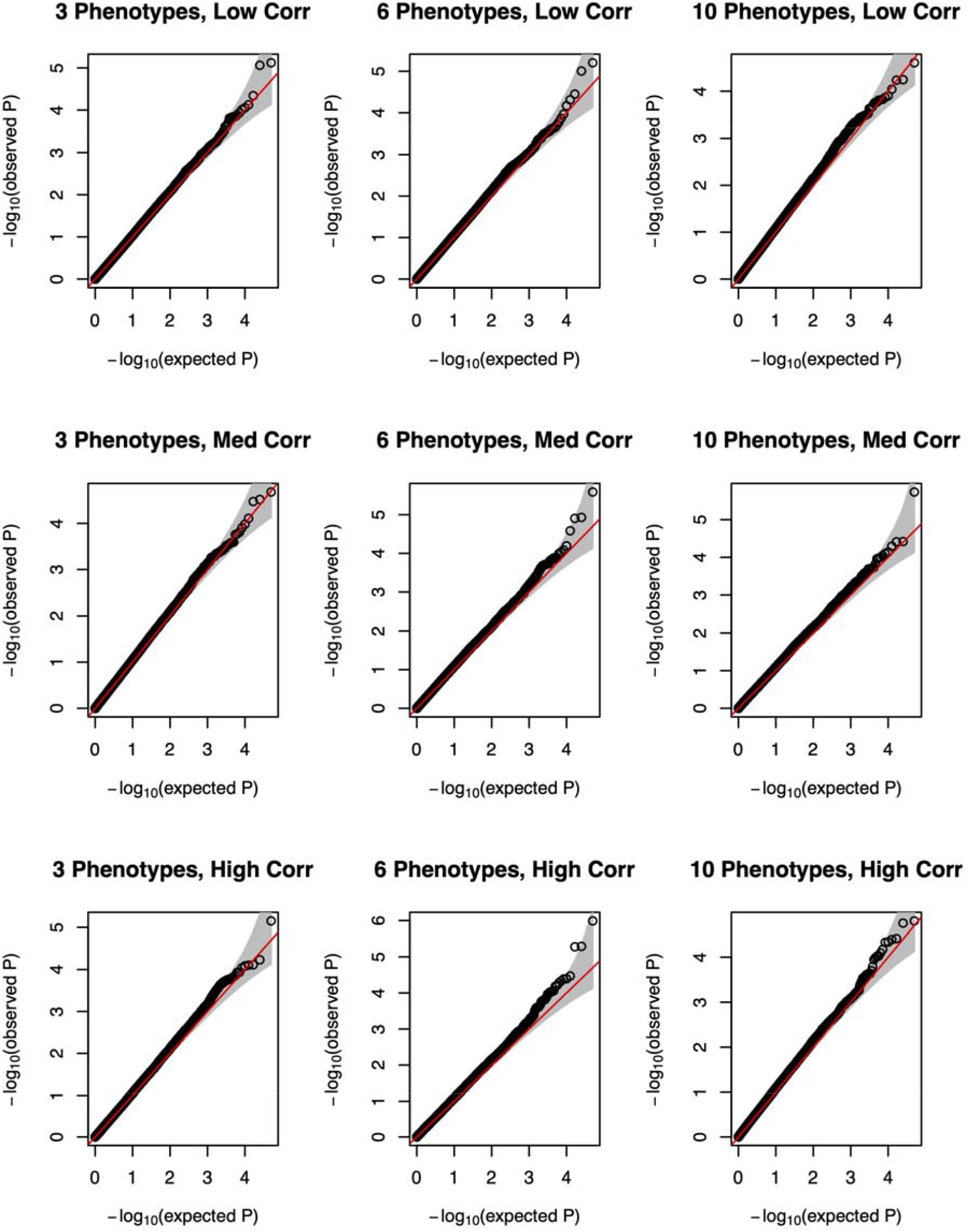
QQ plots of p-values for proposed parent-of-origin effect test under the null hypothesis using a series of 50,000 simulations of 5,000 individuals using 3 (left column), 6 (middle column) or 10 (right column) continuous non-normal phenotypes. MAF is assumed to be 0.25. Horizontal panels depict level of pairwise-trait correlation (low, medium, high). Abbreviations: QQ, quantile-quantile; MAF, minor allele frequency.

### 3.2 Power

Simulation results comparing the performance of POIROT to the competing univariate test under the assumption of true POE(s) are summarized in Figure 3. This figure reflects normally distributed traits and sample size of 5,000 (*α* = 5 X 10^−4^). Corresponding results from all other additional power settings, including both normal and non-normal traits, sample sizes of 5,000 and 10,000, and *α* = 0.005, 5 X 10^−4^ are provided in Supplemental Figures 2-9.

**Figure 3.**
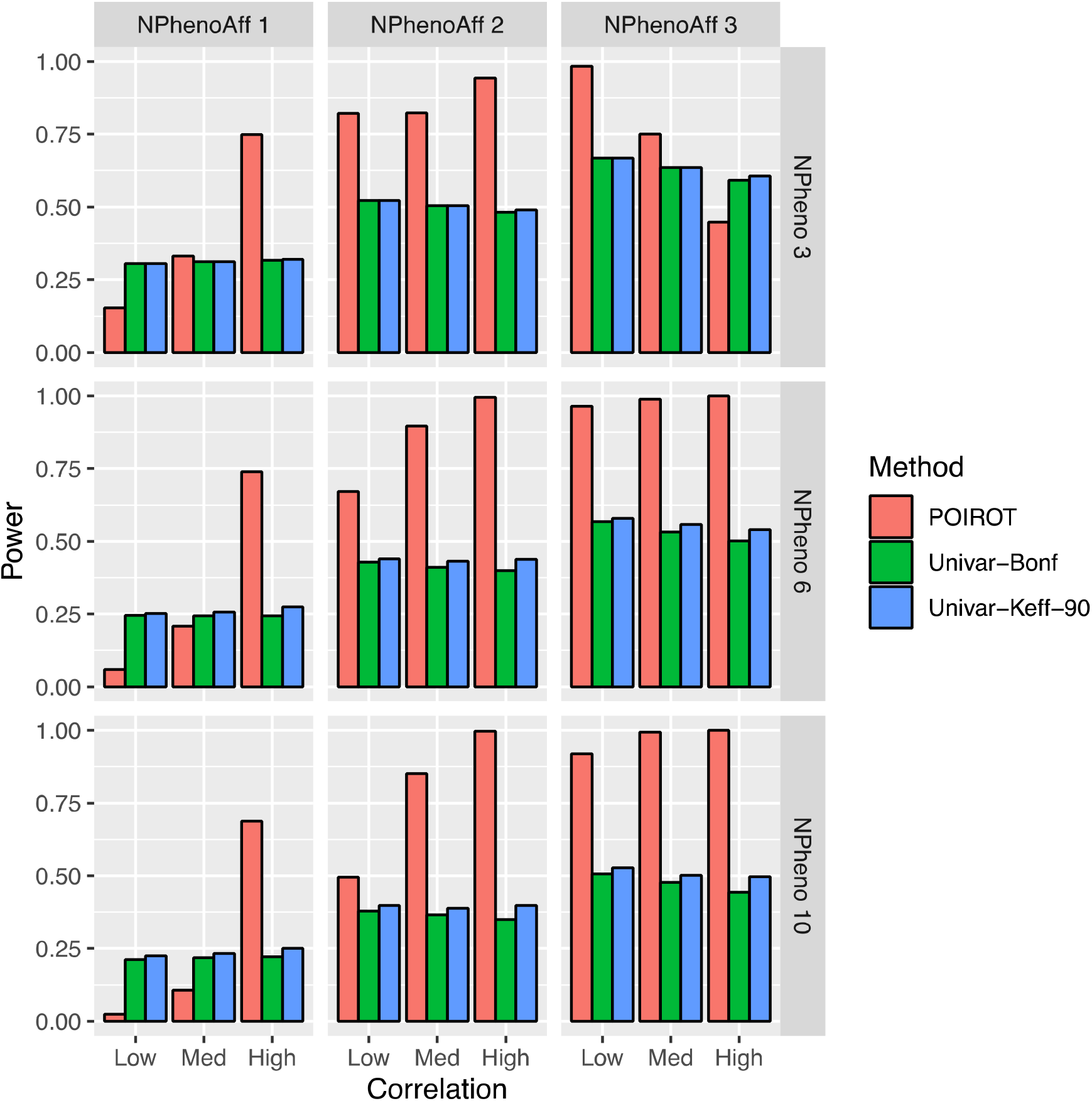
Power of POIROT to identify POEs assuming *K* = 3, 6, or 10 normal phenotypes (horizontal panels) compared to univariate test. We assume either 1, 2, or 3 of the phenotypes harbor POEs at the locus (vertical panels). We performed 5,000 simulations for each scenario. We calculated power at significance level 0.0005 for our multi-trait test and 0.0005/*K* (Bonferroni correction) and 0.0005/*K*_*eff*_ for the univariate test, where *K*_*eff*_ is the number of PCs needed to explain 90% phenotypic variation. for POE traits, MAF = 0.25, and sample size = 5,000. Abbreviations: POE, parent-of-origin effect; MAF, minor allele frequency; PCs, principal components.

Simple Bonferroni correction tends to be overly conservative in the presence of correlated traits. We therefore used two multiple-testing correction approaches for the univariate method. As power generally increases with increasing sample size and POE magnitude, the scenarios shown in Figure 3 correspond to a β_Mk_ of 0.75 and sample size of 5,000. For almost all scenarios, we see three general trends. First, unlike the univariate method, our method successfully leverages the correlation among phenotypes. We see power increasing with increasing trait correlation. Second, when pleiotropy exists and more than one phenotype harbors a POE, our method outperforms the univariate approach regardless of the multiple testing correction strategy. Third, power of POIROT increases as the number of phenotypes associated with the maternally-transmitted alternative allele increases across all levels of phenotypic correlation.

The one exception to these trends is the top right panel of Figure 3. This reflects the scenario where 3 of 3 phenotypes harbor POEs of the same magnitude and direction. We see here that power decreases going from low to medium correlation and from medium to high correlation. We also see lower power when 3 phenotypes are affected when compared to the corresponding settings when only 2 of 3 phenotypes have POEs. This pattern, although unusual, has been documented in previous cross-phenotype methodological studies (Ray *et al*., 2016; Broadaway *et al*., 2016). As described in Section 2.2, the R-Omnibus test for equality of covariance matrices used by POIROT ultimately employs a one-way MANOVA test to generate our test statistic. Ray et al. describe how when we have *K* correlated traits being tested and a SNP is associated with all *K* traits, utilizing a MANOVA to find marginal associations with multiple traits can result in an appreciable loss of power. In particular, the authors show how the power of MANOVA is asymptotically lower when all traits are associated with equal magnitude and direction than when fewer than *K* phenotypes are associated (Ray *et al*., 2016).

### 3.3 Applied Data Analysis

We applied our method for detecting POEs to genotype and multivariate phenotype data of 1,289 unrelated infants of European ancestry from the Hyperglycemia and Adverse Pregnancy Outcome (HAPO) Study. Raw phenotype measures were quantitative anthropometric measures related to infant size at birth (birth weight, length, head circumference, and three skinfold measurements). Phenotypes were appropriately adjusted for the effects of the first two PCs, infant sex, gestational age at birth, maternal BMI, and maternal smoking frequency. For the 6,219,272 variants considered, the average computation time per test was 0.58 seconds. Analysis was run with parallel computation, and time per chromosome ranged between 8.4 and 117.3 hours (median 39.0 hours). In short, although we did not see any variants falling below the Bonferroni-adjusted genome-wide significance threshold of 5 × 10^−8^, we saw one SNP with near genome-wide significance (rs1496904, POIROT *p* = 9.58 × 10^−8^). This SNP is 138kb from the transcription start site of gene *SEMA6D*. Common polymorphisms in this gene have previously been associated with arm fat mass, leg fat mass, body fat percentage, height, and other adult-correlates of traits similar to those we tested in the HAPO study infants (http://www.nealelab.is/uk-biobank/, Ochoa *et al*., 2021; Kichaev *et al*., 2019)). Thirteen other variants at this locus (chr15:47321206-47355147) similarly had POIROT p-values below 5 × 10^−7^. As we see in the Manhattan plot of Figure 4b, there is another locus of suggestive significance on chromosome 1 (chr1:154328785-154347720) with six variants whose p-values fall below 5 × 10^−7^. The lead SNP is rs141140594 (POIROT *p* = 2.43 × 10^−7^). This SNP lies 3kb from gene *ATP8B2*. Nearby variants have previously been associated with type 2 diabetes. However, the mechanisms by which this gene is functionally implicated in the disease remain unclear (Imamura *et al*., 2016; M *et al*., 2020). Furthermore, these loci were not identified by the univariate approach across the six tests for each phenotype (minimum *p* = 5.42 × 10^−5^).

**Figure 4.**
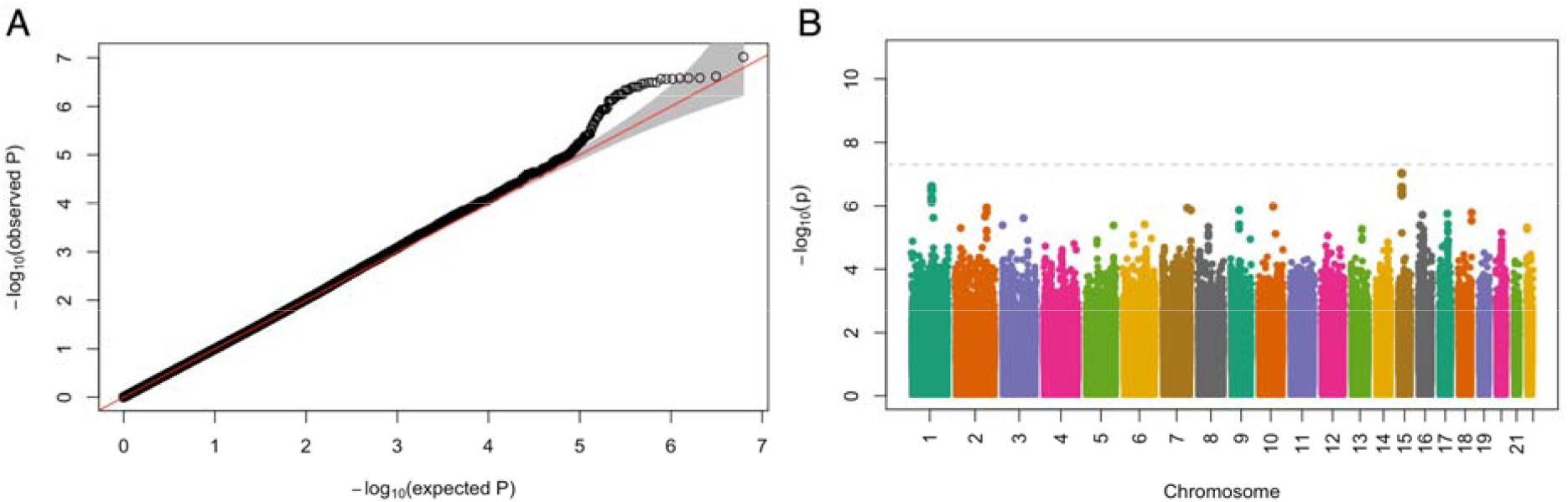
A) QQ plot from parent-of-origin effects analysis using POIROT and six HAPO study phenotypes related to infant size at birth. B) Manhattan plot of corresponding analysis where dashed line represents genome-wide significance of . Abbreviations: QQ, quantile-quantile.

## 4 DISCUSSION

In this paper, we introduce a multivariate method, POIROT, for identifying common variants exhibiting POEs on one or more quantitative phenotypes in unrelated subjects. This work is motivated dually by the widespread evidence of pleiotropy in the genetics literature, as well as the limited statistical options for detecting POEs in unrelated cohorts. Our proposed method is an inherently simple statistical test of whether the phenotypic covariance matrix of heterozygotes is equal to that of homozygotes at a given locus. It represents a multivariate extension of the POE test of a single continuous phenotype proposed by Hoggart et al. (Hoggart *et al*., 2014). It allows for appropriate adjustment for the effects of important covariates on the phenotypes under study and is also computationally efficient for application to biobank-scale datasets (Supplemental Tables 2-3). The R code for implementing POIROT is publicly available (see Data Availability).

Through simulations, we demonstrate POIROT achieves appropriate type I error under the null. It further displays superior power to detect POEs than the competing univariate approach under most settings. Our method is indeed robust to non-normality of phenotypes across several simulation scenarios. We further applied our method to real GWAS data on unrelated infants from the HAPO Study. In this analysis, we considered six anthropometric measurements at birth related to fetal growth. Although the analysis presented here did not reveal any variants meeting the stringent genome-wide significance threshold, two loci of suggestive significance were identified that may warrant further investigation. These loci are not located within 500kb of any known imprinting gene in humans. They may, however, be strong candidates for follow-up replication analyses using independent trio studies or other familial studies of these phenotypes.

The top locus has been shown in prior studies to be associated directly with similar adult anthropometric measures. Further, the second has documented associations with type 2 diabetes, a condition of the metabolic syndrome. The Barker hypothesis posits that inadequate fetal nutrition, quantitative measures of which include birth weight, confers greater risk of metabolic syndrome later in life (Edwards, 2017). We also note that these loci were not identified by the competing univariate approach. This suggests that joint consideration of multiple related traits can indeed help improve discovery of POE variants. We do note that such discovery potential is limited in the HAPO dataset due to sample size (N=1,289). This dataset is small compared to many modern consortium GWAS and is vastly unpowered to detect even marginal effects of SNPs affecting body size or type 2 diabetes (Xue *et al*., 2018; Berndt *et al*., 2013). In our simulations, we show significant power in sample sizes 5-10 times larger than that of the HAPO analysis. That any plausible suggestive results are observed by our method in this dataset, we take as a promising sign for future work.

There are several avenues we are interested in pursuing to extend the work presented here. Rather than testing genome-wide variants, implementation of a two-stage screening procedure may mitigate the multiple testing burden. In the first stage, we propose to perform a standard GWAS for marginal (not parent-of-origin dependent) variant associations that considers multiple traits jointly. We restrict consideration to marginal association tests that are orthogonal to POIROT and thus provide complementary information. We can then efficiently test a smaller subset of top SNPs identified from the first stage for POEs. Another limitation we acknowledge is the requirement of continuous phenotypes. We are interested in the possible extension of our approach to accommodate dichotomous multivariate traits. One potential solution would be to use liability-threshold models (Hujoel *et al*., 2020) that can effectively transform a binary outcome into a continuous-valued posterior mean genetic liability.

## ACKNOWLEDGEMENTS

The data used for the applied analysis described in this paper were downloaded from the database of Genotypes and Phenotypes (dbGaP, http://www.ncbi.nlm.nih.gov/gap) with study accession phs000096.v4.p1. The Hyperglycemia and Adverse Pregnancy Outcome (HAPO) Study was a part of the Gene Environment Association (GENEVA) studies, and we acknowledge the principal investigator William L. Lowe, all co-investigators, the National Human Genome Research Institute (NHGRI), and NIH grant U01 HG004415.

## Data availability

The code for implementing this method in R is publicly available at https://github.com/staylorhead/POIROT-POE.

