## Supplemental Figures for "POIROT: A powerful test for parent-of-origin effects in unrelated samples leveraging multiple phenotypes"

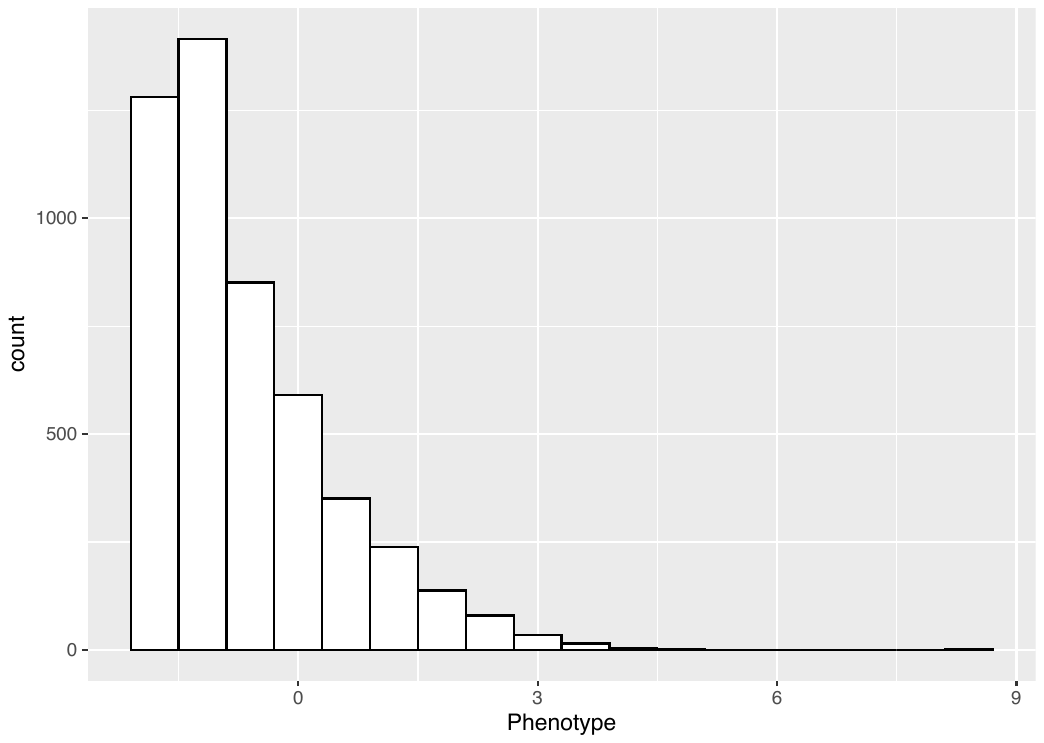

**Supplemental Figure 1.** Histogram of example simulated non-normal phenotypes assuming skewness = 2 and excess kurtosis = 2. Data shown here corresponds to a sample size of 5,000 for a single phenotype with no parent-of-origin effects.

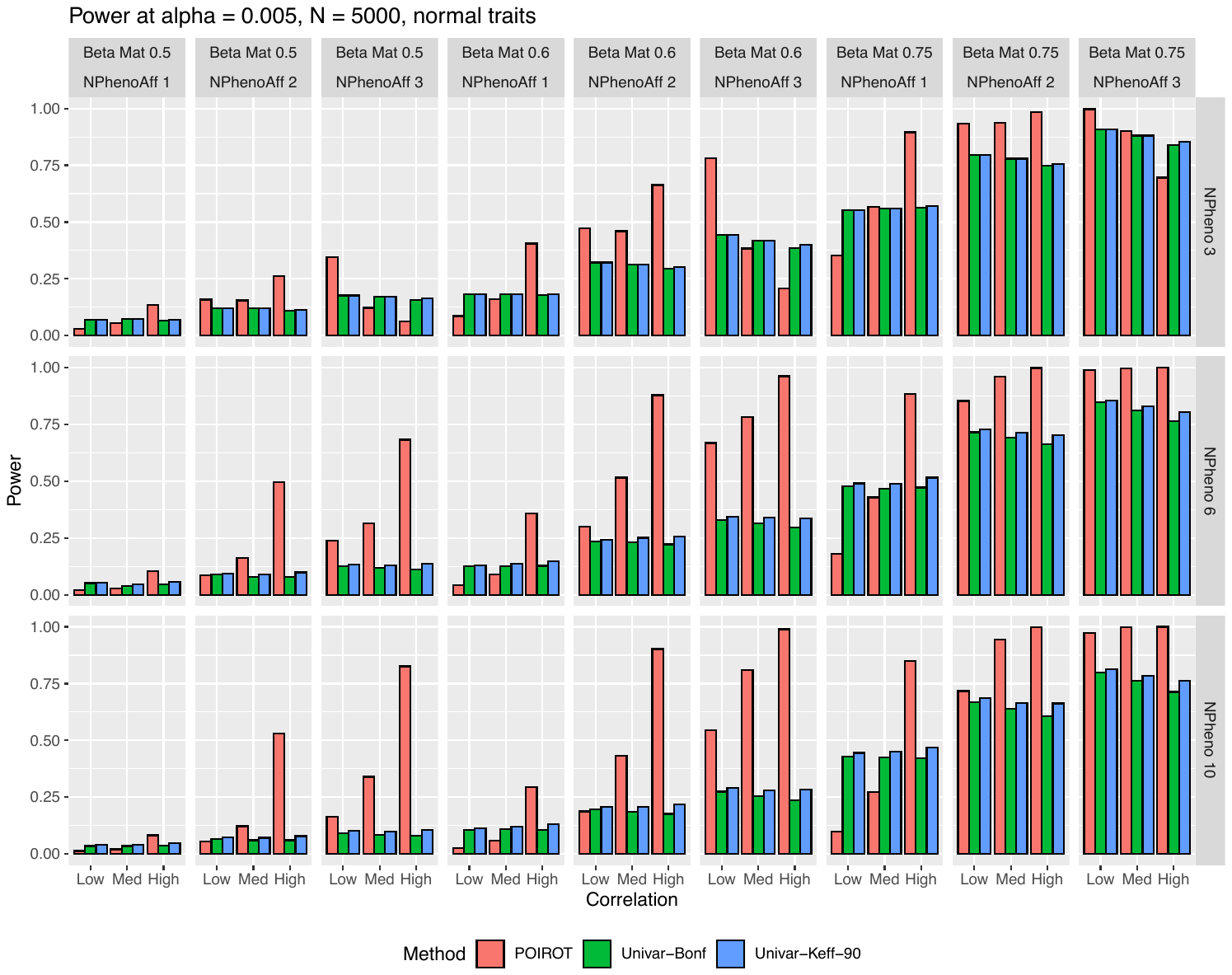

**Supplemental Figure 2.** Power of POIROT to identify POEs assuming *K* = 3, 6, or 10 normal phenotypes (horizontal panels) compared to univariate test. We assume either 1, 2, or 3 of the phenotypes harbor POEs at the locus with varying magnitude of $\beta_{Mk}$ (vertical panels). We performed 5,000 simulations for each scenario. We calculated power at significance level 0.005 for our multi-trait test and 0.005/*K* (Bonferroni correction) and 0.005/*K_eff_* for the univariate test, where *K_eff_* is the number of PCs needed to explain 90% phenotypic variation. We assume MAF = 0.25 and sample size = 5,000. Abbreviations: POE, parent-of-origin effect; MAF, minor allele frequency; PCs, principal components.

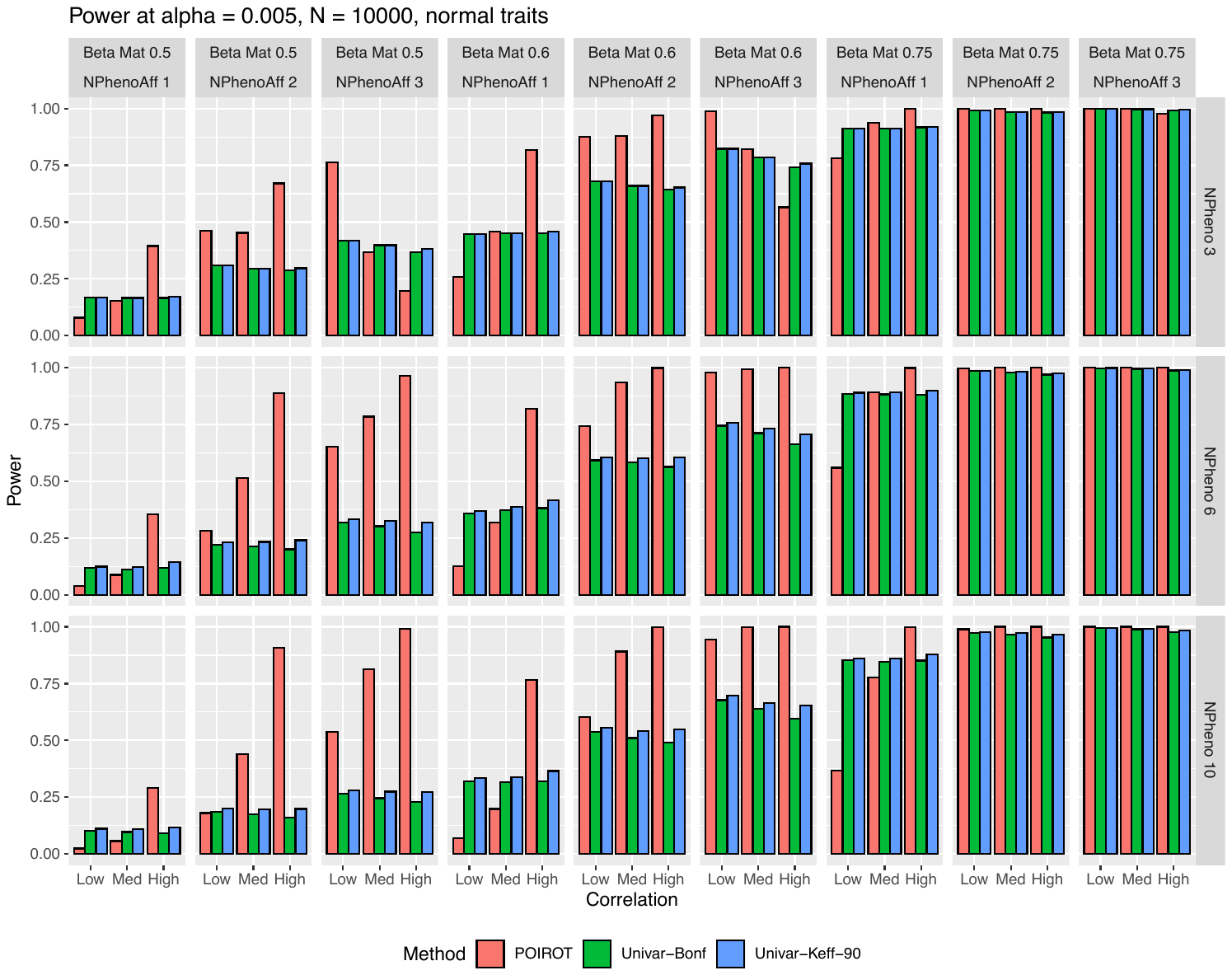

**Supplemental Figure 3.** Power of POIROT to identify POEs assuming *K* = 3, 6, or 10 normal phenotypes (horizontal panels) compared to univariate test. We assume either 1, 2, or 3 of the phenotypes harbor POEs at the locus with varying magnitude of $\beta_{Mk}$ (vertical panels). We performed 5,000 simulations for each scenario. We calculated power at significance level 0.005 for our multi-trait test and 0.005/*K* (Bonferroni correction) and 0.005/*K_eff_* for the univariate test, where *K_eff_* is the number of PCs needed to explain 90% phenotypic variation. We assume MAF = 0.25 and sample size = 10,000. Abbreviations: POE, parent-of-origin effect; MAF, minor allele frequency; PCs, principal components.

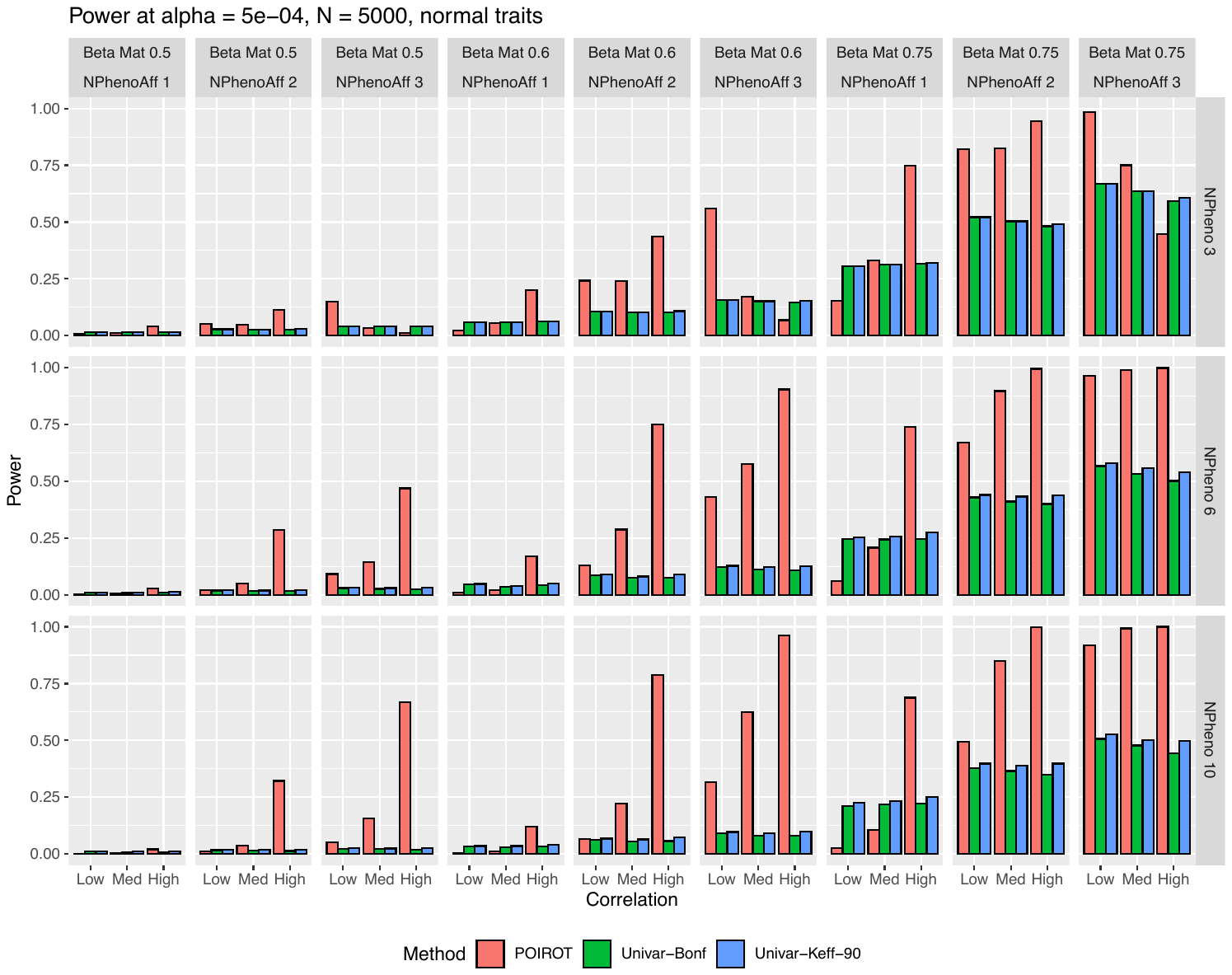

**Supplemental Figure 4.** Power of POIROT to identify POEs assuming *K* = 3, 6, or 10 normal phenotypes (horizontal panels) compared to univariate test. We assume either 1, 2, or 3 of the phenotypes harbor POEs at the locus with varying magnitude of $\beta_{Mk}$ (vertical panels). We performed 5,000 simulations for each scenario. We calculated power at significance level 5×10^-4^ for our multi-trait test and 5×10^-4^/*K* (Bonferroni correction) and 5×10^-4^/*K_eff_* for the univariate test, where *K_eff_* is the number of PCs needed to explain 90% phenotypic variation. We assume MAF = 0.25 and sample size = 5,000. Abbreviations: POE, parent-of-origin effect; MAF, minor allele frequency; PCs, principal components.

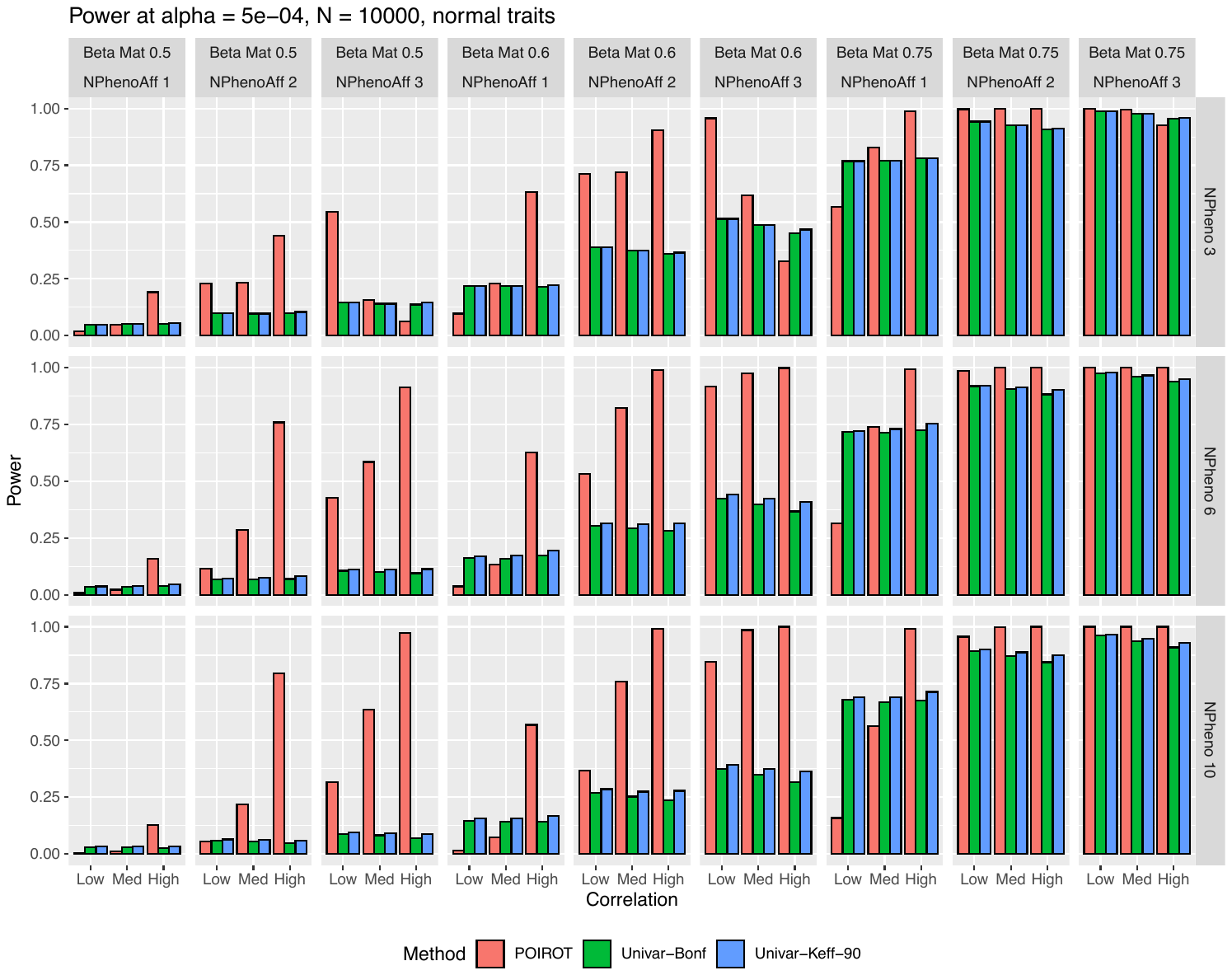

**Supplemental Figure 5.** Power of POIROT to identify POEs assuming *K* = 3, 6, or 10 normal phenotypes (horizontal panels) compared to univariate test. We assume either 1, 2, or 3 of the phenotypes harbor POEs at the locus with varying magnitude of $\beta_{Mk}$ (vertical panels). We performed 5,000 simulations for each scenario. We calculated power at significance level 5×10^-4^ for our multi-trait test and 5×10^-4^/*K* (Bonferroni correction) and 5×10^-4^/*K_eff_* for the univariate test, where *K_eff_* is the number of PCs needed to explain 90% phenotypic variation. We assume MAF = 0.25 and sample size = 10,000. Abbreviations: POE, parent-of-origin effect; MAF, minor allele frequency; PCs, principal components.

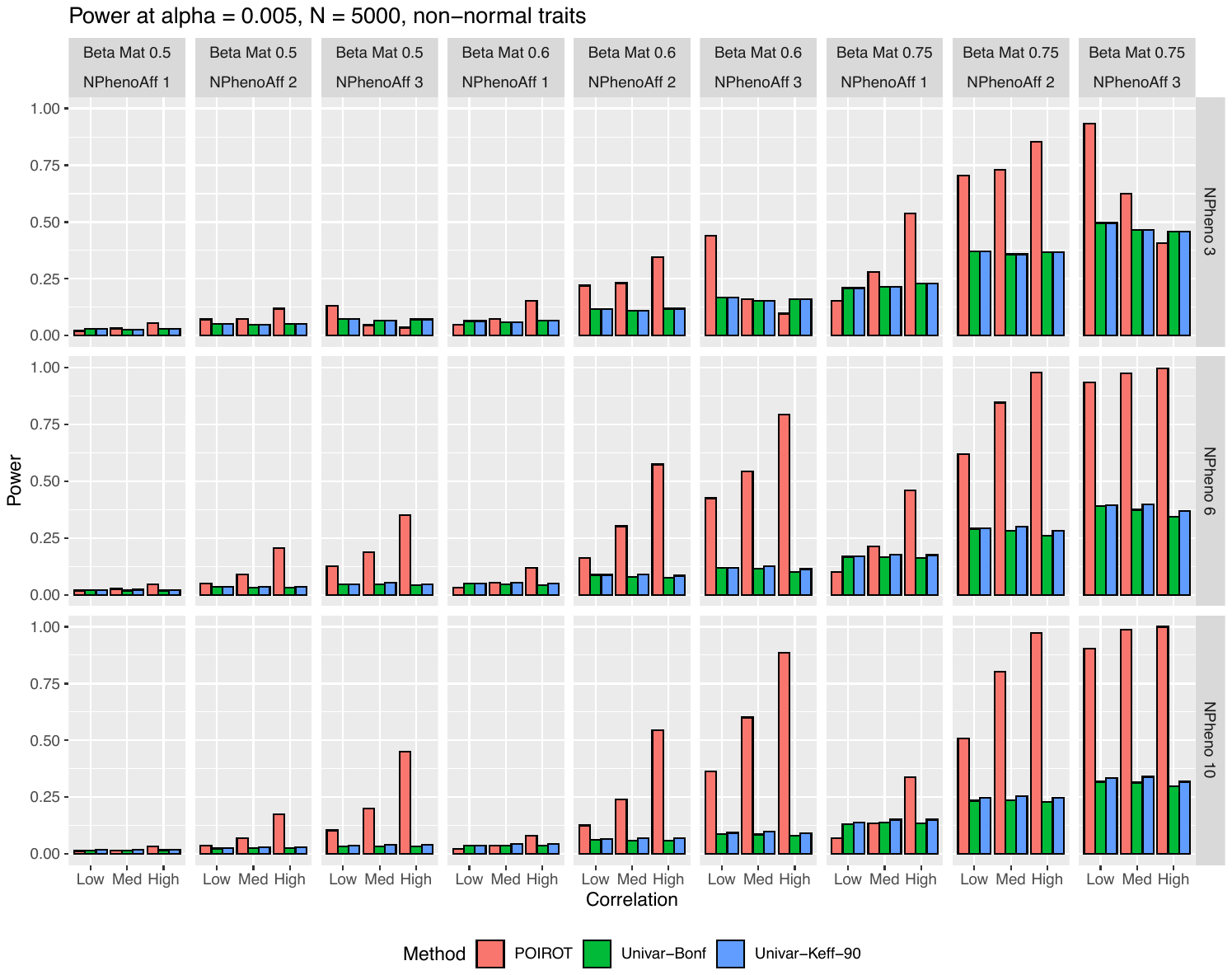

**Supplemental Figure 6.** Power of POIROT to identify POEs assuming *K* = 3, 6, or 10 non-normal phenotypes (horizontal panels) compared to univariate test. We assume either 1, 2, or 3 of the phenotypes harbor POEs at the locus with varying magnitude of $\beta_{Mk}$ (vertical panels). We performed 5,000 simulations for each scenario. We calculated power at significance level 0.005 for our multi-trait test and 0.005/*K* (Bonferroni correction) and 0.005/*K_eff_* for the univariate test, where *K_eff_* is the number of PCs needed to explain 90% phenotypic variation. We assume MAF = 0.25 and sample size = 5,000. Abbreviations: POE, parent-of-origin effect; MAF, minor allele frequency; PCs, principal components.

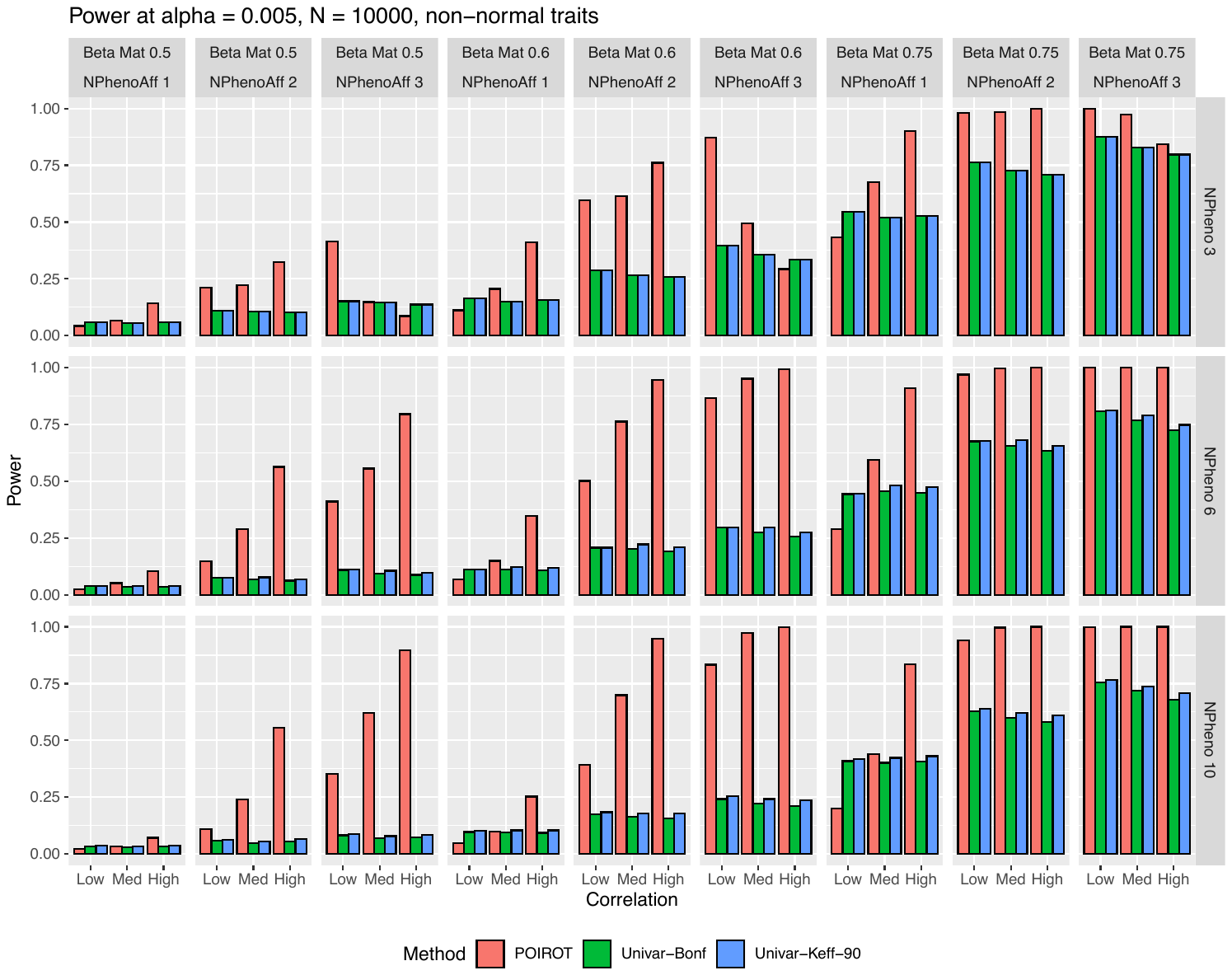

**Supplemental Figure 7.** Power of POIROT to identify POEs assuming *K* = 3, 6, or 10 non-normal phenotypes (horizontal panels) compared to univariate test. We assume either 1, 2, or 3 of the phenotypes harbor POEs at the locus with varying magnitude of $\beta_{Mk}$ (vertical panels). We performed 5,000 simulations for each scenario. We calculated power at significance level 0.005 for our multi-trait test and 0.005/*K* (Bonferroni correction) and 0.005/*K_eff_* for the univariate test, where *K_eff_* is the number of PCs needed to explain 90% phenotypic variation. We assume MAF = 0.25 and sample size = 10,000. Abbreviations: POE, parent-of-origin effect; MAF, minor allele frequency; PCs, principal components.

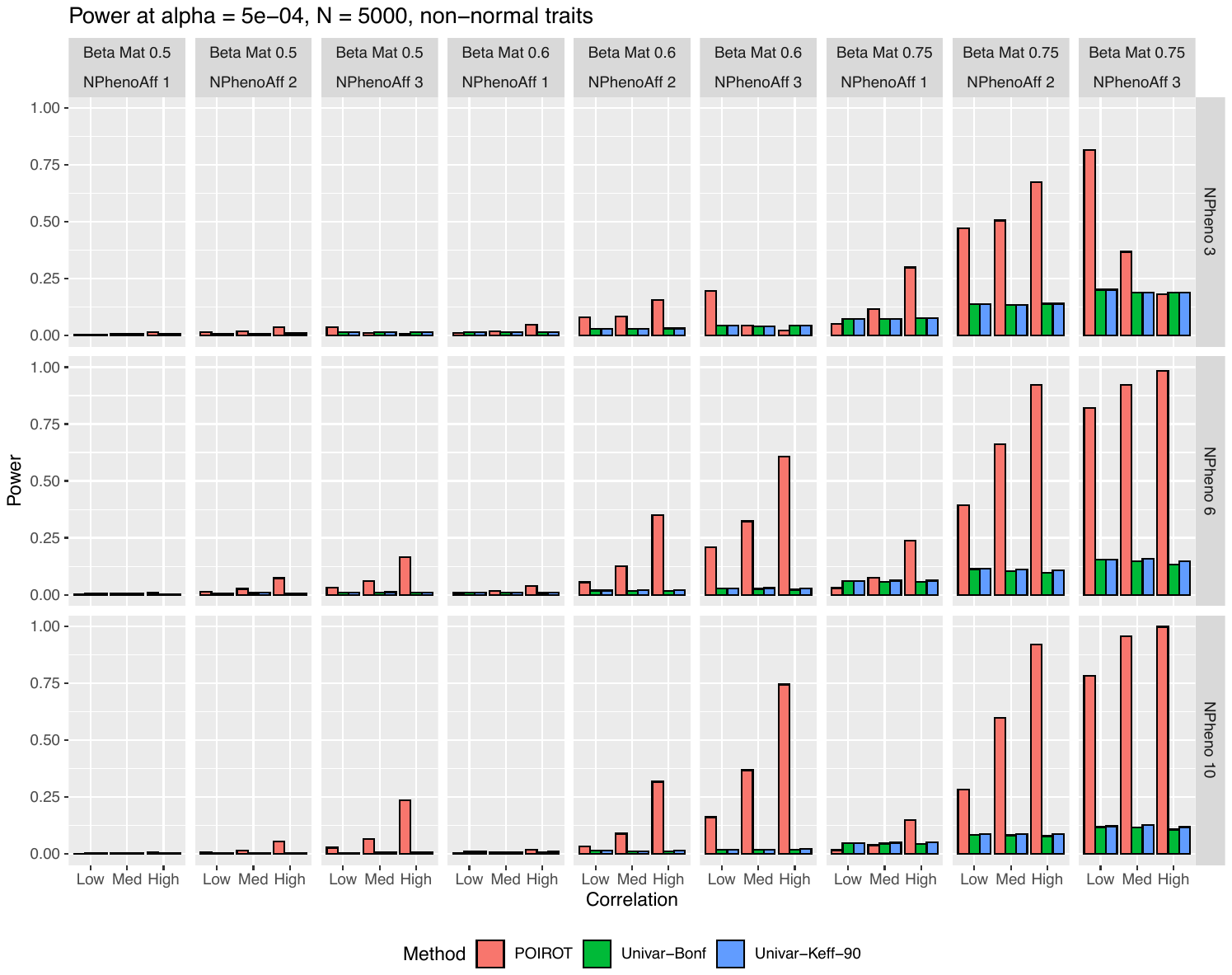

**Supplemental Figure 8.** Power of POIROT to identify POEs assuming *K* = 3, 6, or 10 non-normal phenotypes (horizontal panels) compared to univariate test. We assume either 1, 2, or 3 of the phenotypes harbor POEs at the locus with varying magnitude of $\beta_{Mk}$ (vertical panels). We performed 5,000 simulations for each scenario. We calculated power at significance level 5×10^-4^ for our multi-trait test and 5×10^-4^/*K* (Bonferroni correction) and 5×10^-4^/*K_eff_* for the univariate test, where *K_eff_* is the number of PCs needed to explain 90% phenotypic variation. We assume MAF = 0.25 and sample size = 5,000. Abbreviations: POE, parent-of-origin effect; MAF, minor allele frequency; PCs, principal components.

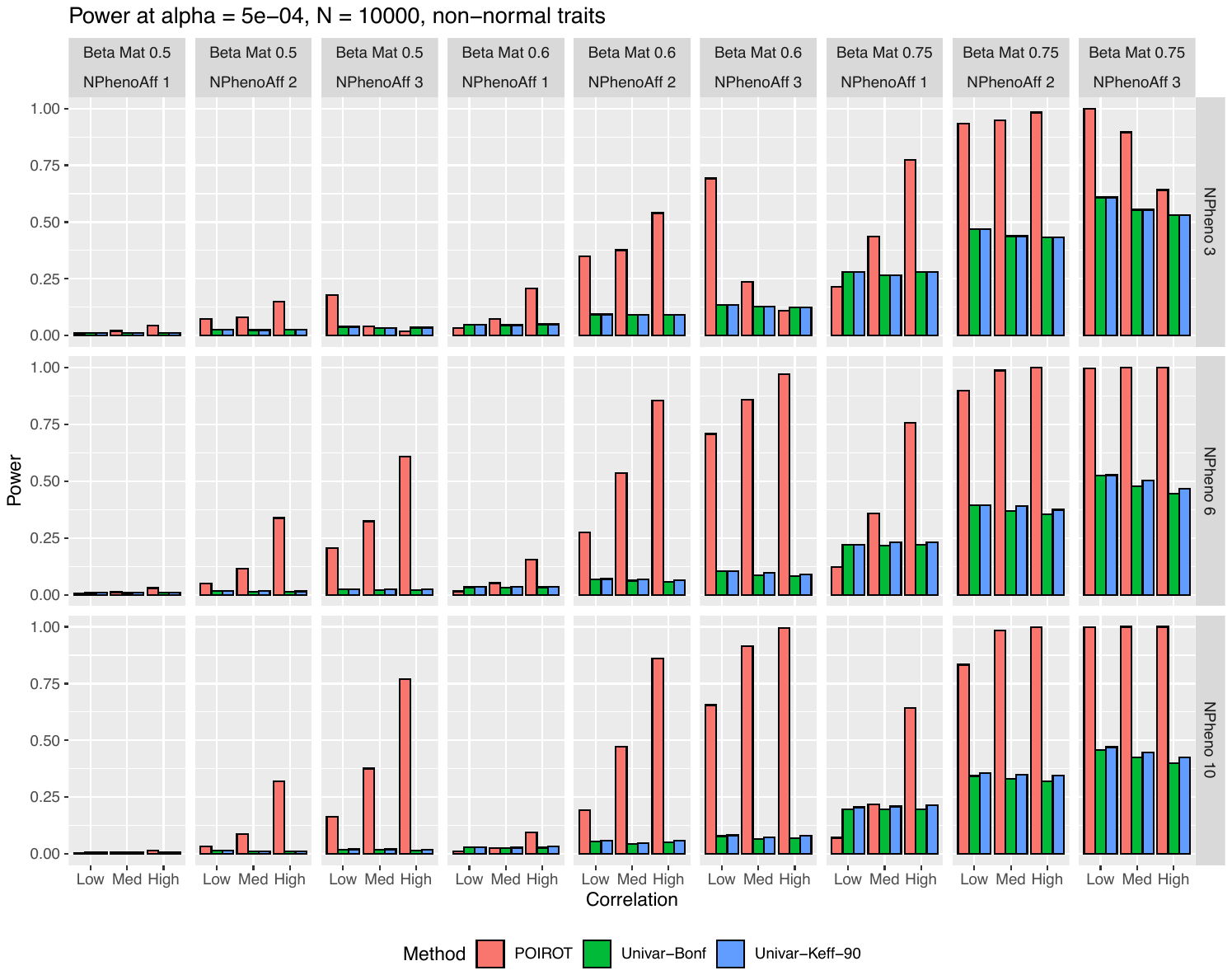

**Supplemental Figure 9.** Power of POIROT to identify POEs assuming *K* = 3, 6, or 10 non-normal phenotypes (horizontal panels) compared to univariate test. We assume either 1, 2, or 3 of the phenotypes harbor POEs at the locus with varying magnitude of $\beta_{Mk}$ (vertical panels). We performed 5,000 simulations for each scenario. We calculated power at significance level 5×10^-4^ for our multi-trait test and 5×10^-4^/*K* (Bonferroni correction) and 5×10^-4^/*K_eff_* for the univariate test, where *K_eff_* is the number of PCs needed to explain 90% phenotypic variation. We assume MAF = 0.25 and sample size = 10,000. Abbreviations: POE, parent-of-origin effect; MAF, minor allele frequency; PCs, principal components.
